# A Tale of Two Cell Lines: Characterization of differential efficacy of small molecule drugs cediranib and NU-7441 on primary vs. metastatic colorectal cancer

**DOI:** 10.1101/2025.09.18.677116

**Authors:** Josephine D. Tsang, Cai M. Roberts

## Abstract

Colorectal cancer is recognized as one of the leading causes of cancer death amongst both sexes in the U.S. Despite rigorous screening, those diagnosed with colon cancer often face poor prognosis, and approximately 70% of affected patients will develop metastatic relapse. The investigation of colon carcinoma cell lines’ genetic variability and response to chemotherapy panels may aid in targeting therapies to improve outcomes. This study aims to find correlations between metastasis status, gene variability, and drug response. We used two cell lines that were isolated from the same 51-year-old male with colorectal adenocarcinoma: a primary tumor-derived line (SW480) and a secondary metastasis-derived line (SW620). Live cell imaging using time-lapse microscopy over three days exhibited differential cell death responses following treatment with multiple chemotherapeutic agents, particularly cediranib and Nu-7441, with SW620 demonstrating greater sensitivity. Western blots revealed changes in DNA repair machinery expression (particularly NHEJ proteins) between SW480 and SW620. RNA sequencing and Gene Ontology analysis corroborated our findings, demonstrating upregulated DNA repair and metabolic survival genes including TGM2 in SW620 and PROM1 in SW480. An SW620 line grown in non-attachment plates and then reattached (SW620F) exhibited high DNA-PKcs activation and drug sensitivity. Correlation between drug response and gene expression crucial to cell growth and successful metastasis may reveal new biomarkers to target potential treatments.

## 1 INTRODUCTION

Colorectal cancer (CRC) is one of the leading causes of cancer morbidity and mortality amongst both sexes in the United States [1] . Those with a higher risk of CRC, such as those with a family history or those with a genetic predisposition often receive colon cancer screening at a younger age to identify malignant polyps. However, despite rigorous screening, colon cancer remains a prevalent cause of cancer death. Metastasis is common, and 40% of patients present with relapsed disease [1] . The five-year survival rate for CRC is approximately 63%, but drops to 13% for metastatic CRC [2] . The main difficulty in identifying and treating colorectal cancer is attributed to its insidious presentation, propensity for metastasis, and drug resistance.

Current therapies targeted at CRC include oxaliplatin, irinotecan, and capecitabine [1] . However there has been difficulty in characterizing specific curative models for treating metastatic CRC due to the inherent limitations of chemotherapeutic treatments. Patients often experience systemic toxicity, low tumor specificity, and unpredictable innate and acquired resistance. Our research is aimed at identifying molecular and pathological features to guide the selection of chemotherapeutic treatment. Previous research in epithelial to mesenchymal transition (EMT) in ovarian and breast cancer was used as a model to guide experimentation. EMT involves alterations to DNA repair pathways, which may lead to vulnerability to pharmaceutical interventions targeting those pathways [3] . Double-strand breaks (DSBs) may be repaired either by homologous recombination (HR) or non-homologous end joining (NHEJ). BRCA1 and BRCA2 are essential for HR; BRCA1 is responsible for recognizing and signaling DNA damage, while BRCA2 mediated the loading of the protein Rad51 onto single stranded DNA. Those who express BRCA1 or BRCA2 mutations are at a higher risk of developing breast or ovarian cancer, and may be at higher risk of other cancers, including CRC. Recent studies have identified an increased risk of CRC with BCRA1 and BRCA2 carriers [4] . Rad51 is a recombinase enzyme that plays a central role in HR, working with BRCA1 and BRCA2 to promote the pairing of broken DNA repair strands with homologous sequences. Overexpression of Rad51 has a wide variety of consequences ranging from increased HR and increased resistance to DNA damaging agents to disruption of the cell cycle [5] .

Ku-70 and Ku-80 together form the Ku heterodimer that binds to DSBs, recruit other repair proteins, and stabilize the ends of DNA for NHEJ. Previous studies in CRC have shown a statistically significant association in the binding activity and protein level of Ku-70 and Ku-80. Cytoplasmic protein expression was found in pathological samples, but not in normal tissues [6] . DNA phosphorylase kinase (DNA-PK) is part of the complex of NHEJ with Ku-70 and Ku-80. DNA-PKcs is the catalytic subunit of this enzyme complex.

EMT has also been associated with vascular endothelial growth factor (VEGF), a potent inducer of growth of new blood vessels, also known as angiogenesis, by binding to receptors on endothelial cells. VEGF overexpression is common in tumors and contributes to their growth and survival by promoting angiogenesis. Both fibronectin and VEGF contribute to creating a microenvironment that promotes tumor growth and survival by maximizing cellular adhesion and viability.

This study aims to find correlation between metastasis status, gene variability, and drug response in CRC lines. Therapy aimed at specific cellular responses, such as DNA repair in metastatic cells, may reduce relapse and improve survival.

## 2 MATERIALS AND METHODS

### 2.1 Cell lines and culture conditions

Primary colorectal cell line SW480 and secondary metastatic SW620 were isolated from the same patient diagnosed with colorectal adenocarcinoma. Cells were purchased from ATCC (Manassas, VA). SW620F was generated by growing SW620 cells for 1 week in ultra-low attachment plates and then returning them to standard culture vessels. All lines were maintained in DMEM with 10% FBS and 1% penicillin-streptomycin in a 37°C 5% CO_2_ incubator.

### 2.2 RNA sequencing and Gene Ontology

RNA sequencing (Figure 1A) was used to quantify, profile, and analyze the complete transcription of cells. RNA sequencing provides a global view of gene expression changes between SW480 and SW620, the primary and secondary metastatic lines respectively, enabling further insight into the molecular effects of chemotherapeutic drugs. Cells were cultured under standard conditions and harvested. RNA was extracted using the Total RNA Extraction Kit from IBI Scientific (Dubuque, IA) following the manufacturer’s protocol. RNA sequencing was performed by Genewiz by Azenta. Library preparation consisted of rRNA and DNA depletion. Paired-end reads were obtained for biological triplicates. Gene Ontology (GO) analysis (Figure 1B) was conducted on differentially expressed genes to identify biological processes, molecular functions, and cellular components between SW480 and SW620. Differentially expressed genes were classified into upregulated and downregulated groups, and gene ontology was performed separately for each group. Heatmaps of each gene were generated with hierarchical clustering. Those genes significantly differentially expressed were then inferred to provide insight into potential mechanisms of drug action of each cell line compared to the other.

**FIGURE 1.**
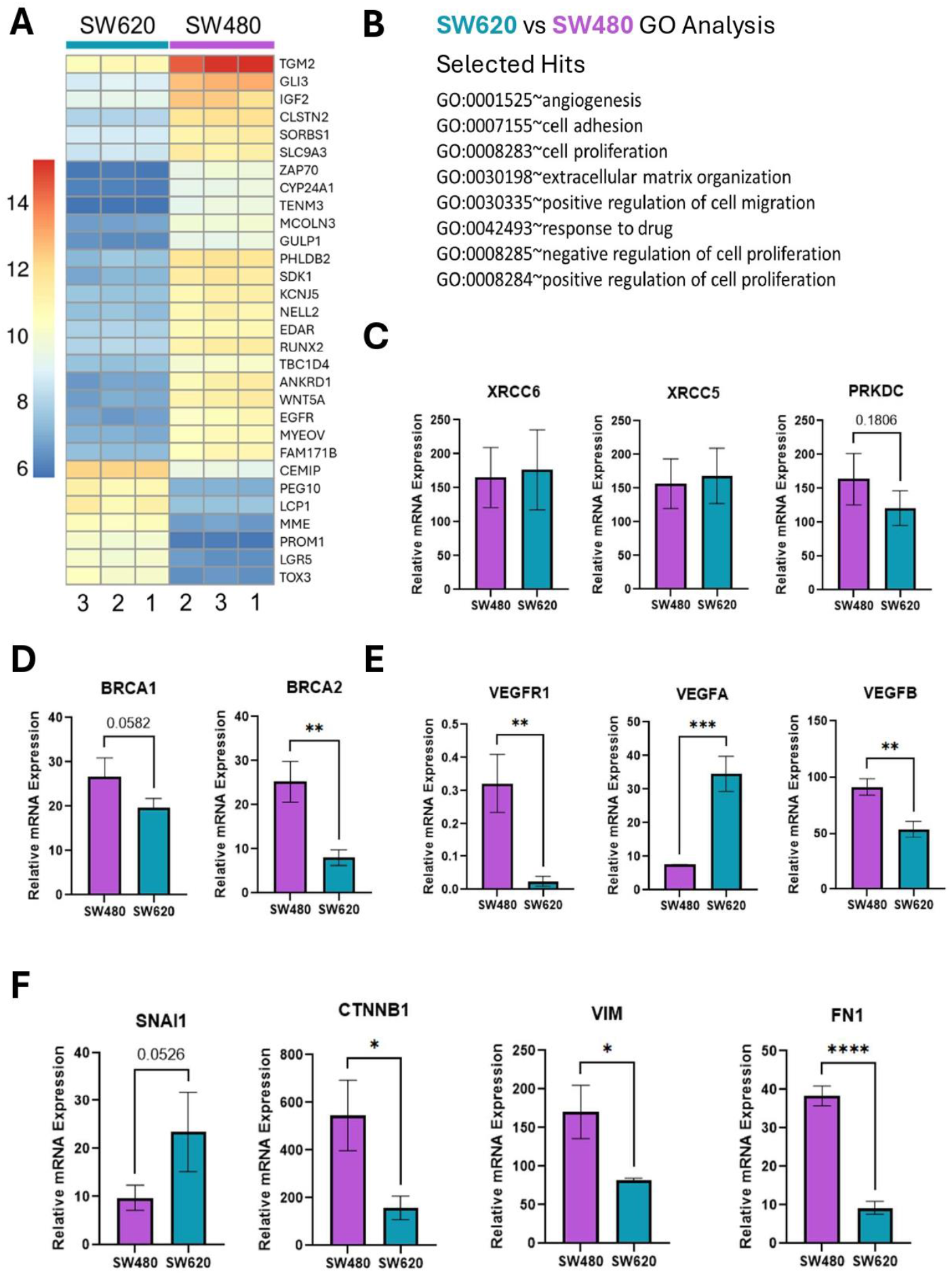
(A) Conditions (genes) between SW480 and SW620 with most significant differences in expression. (B) Selected gene ontology (GO) in the top 20 most statistically significant changed between SW480 and SW620 based on RNAseq. (C) Relative mRNA expression of NHEJ factors XRCC6 (Ku70), XRCC5 (Ku80), and PRKDC (DNA-PKcs). (D) Relative mRNA expression of DNA repair genes BRCA 1 and 2. (E) Relative mRNA expression of angiogenesis factors VEGFR1, VEGFa and VEGFb. (F) Relative mRNA expression of common EMT markers. ^*^p<.05, ^**^p<.01, ^****^p<.0001.

### 2.3 Western Blot

Each cell line was initially analyzed via western blot to identify and validate the proteins responsible for enhancing cell survival. All cells were cultured, harvested, lysed, and analyzed via our published western blot protocol [7] . Primary antibodies were: NHEJ Sampler 76696T, (Cell Signaling, Danvers, MA), BRCA2 (A303-434A, Bethyl Labs, Fortis, Waltham, MA), and Rad51 (H-92, Santa Cruz Biotechnology, Houston, TX). Beta actin (sc-47778, Santa Cruz) was used as endogenous control.

### 2.4 Live Cell Imaging

Once cellular processes were identified, each line was tested against a battery of chemotherapeutic agents. SW480, SW620, and SW620F cells were seeded into a 96 well plate at approximately 7,500 cells per well. Once incubated for 24 hours for adherence and stabilization, each cell line was exposed to chemotherapeutic drugs cediranib, cisplatin, Nu-7441 [8], Ku-55933 [9], and olaparib for 72 hours. The above listed chemotherapeutic agents were chosen based on their effects on DNA repair and NHEJ, as guided by previous research in epithelial to mesenchymal malignancies such as ovarian cancer [3] . Cells were monitored during the incubation period using live cell imaging through the Celloger Mini Plus live cell imaging system from Curiosis (Seoul, Republic of Korea). Brightfield images were captured every 4 hours and analyzed for cell confluency using the Celloger Analysis software.

### 2.5 Statistics and Replication

Pairwise comparisons were done using Student’s two-tailed t-tests. Drug effects were analyzed using one-way ANOVA with multiple comparisons. p<0.05 was considered significant.

## 3 RESULTS

RNA sequencing and GO analysis demonstrated the largest differences in cell adhesion and angiogenesis pathways. Among the most significantly upregulated genes were TGM2 in SW620 and PROM1 in SW480 (Figure 1A). Significant GO terms (Figure 1B) that were altered are remarkable for angiogenesis, cell adhesion, and both up-regulation and down-regulation of cell proliferation. Furthermore, RNA sequencing revealed little difference in NHEJ factors (Figure 1C), a slight drop in BRCA expression (Figure 1D), and differences in VEGF and VEGFR expression in metastatic vs primary cells (Figure 1E). Mesenchymal marker SNAI1 was increased and CTNNB1 reduced in SW620, indicative of EMT. Surprisingly, VIM and FN1 expression were reduced in SW620, though we have previously observed low FN1 levels in advanced oral cancer cells [7] .

To look deeper at DNA repair differences, we performed western blots for HR (BRCA2 and Rad51) and NHEJ proteins (Ku70, Ku80, and DNA-PKcs) (Figure 2). No significant changes in protein expression were seen for BRCA2, with a trend toward elevated Rad51 expression in Sw620 (Figure 2A). There was a trend toward higher expression of Ku80 and far greater DNA-PKcs phosphorylation in SW620 and more so SW620F compared to SW480 cells (Figure 2B).

**FIGURE 2.**
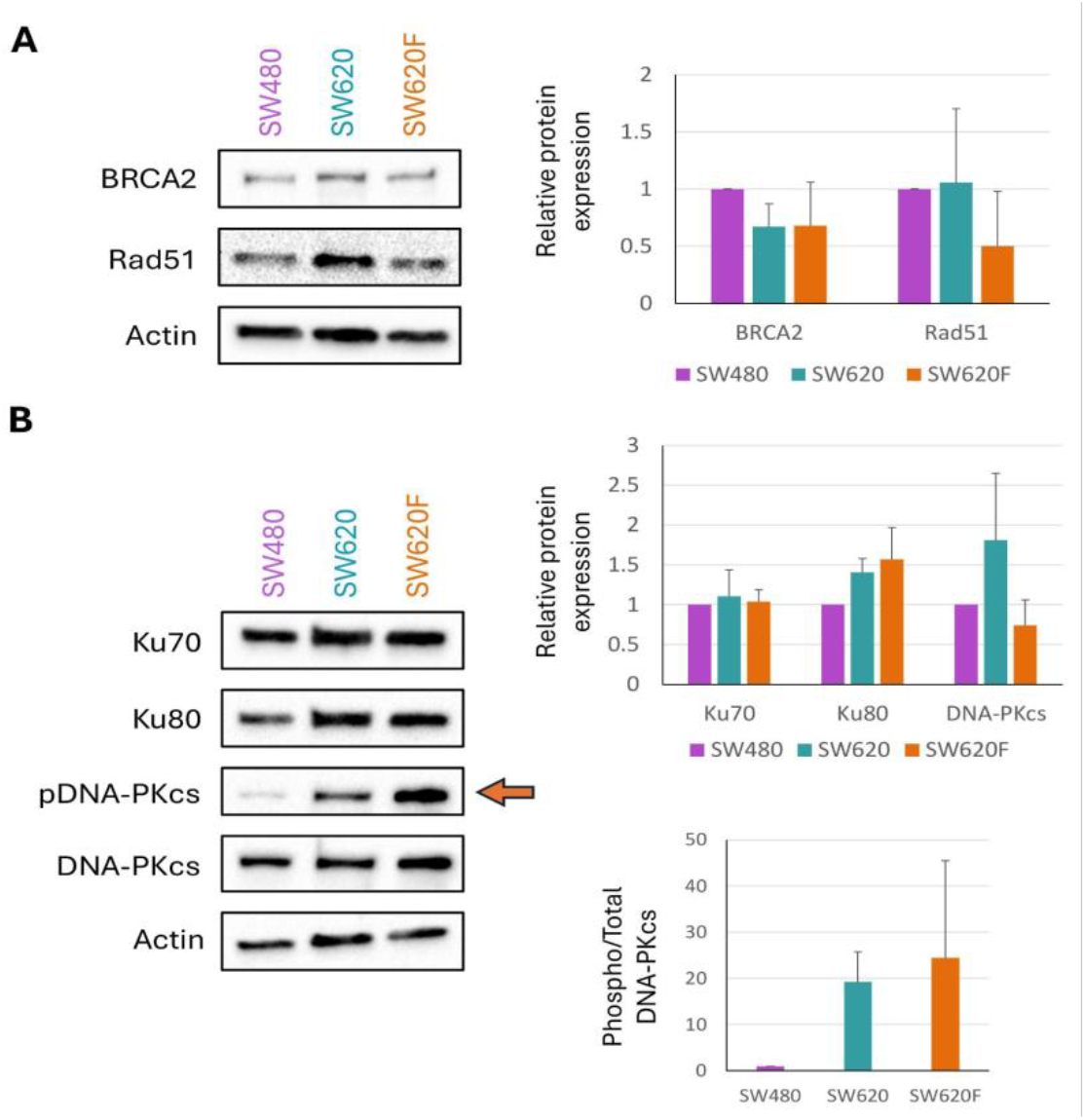
Representative western blots of SW480, SW620, SW620F selected for NHEJ proteins (Ku70, Ku80, DNA-PKcs) and DNA HR proteins (BRCA2 and Rad51), adjacent to proportional depiction of expression. (A) Expression of HR proteins of all 3 cell lines; actin as control. (B) Expression of NHEJ proteins of all 3 cell lines; actin as control. n=3 replicate experiments.

After establishing specific proteins and factors that are involved in cellular survival and tumor metastasis modeling off previous studies in epithelial to mesenchymal transition in ovarian cancer cell lines, SW480 and SW620 were exposed to cisplatin (DNA cross-linker), cediranib (VEGFR inhibitor), Ku-55933 (ATM inhibitor), and Nu-7441 (DNA-PK inhibitor) over a 72 hour period. Live cell imaging revealed the greatest efficacy for cediranib and Nu-55933, while minor differences in growth and survival were noted amongst almost all the chemotherapeutic drugs across the board (Figure 3A). Imaging corroborated findings, and we were able to document hourly changes with each cell line, showing when each cell line begins to plateau. There was demonstrable differential growth when exposed to Cediranib vs. Nu-7441 between SW480, and SW620, and notably SW620F (Figure 3B).

**FIGURE 3.**
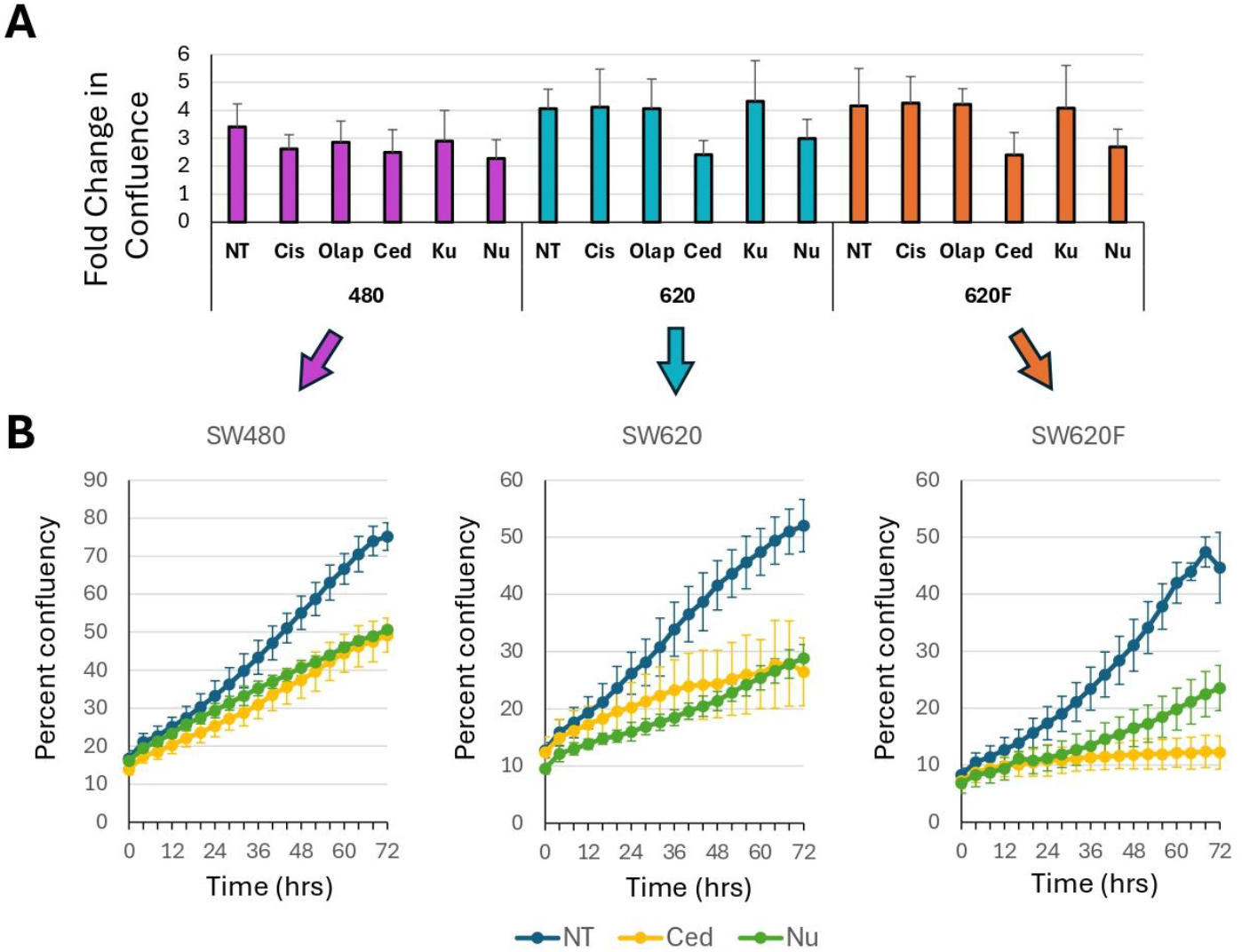
Confluency of chemotherapeutic drugs over a 72 hour period between SW480, SW620, and SW620F. (a) Chemotherapeutic drugs selected depending on its specific target against each cell line. NT, non-treated; Cis, 10µM cisplatin; Olap, 25µM olaparib; Ced, 10µM cediranib; Ku, 10µM Ku-55933; Nu, 10µM Nu-7441. N= 3. (b) Confluency of cells after exposure to chemotherapeutic agents Cediranib and Nu-7441 over time, monitored with live cell imaging. Data from one representative experiment shown.

## 4 DISCUSSION

Colorectal cancer, despite rigorous screening measures, remains one of the most common causes of cancer morbidity and mortality in the United States [1] . Additionally, those in remission from primary CRC have a higher chance of developing metastatic secondary CRC afterward [1] . It is crucial to identify chemotherapeutic agents that target the cellular mechanisms, such as HR and NHEJ, to pinpoint either primary or secondary metastatic CRC. Using targeted therapy eliminates the need for unnecessary trials of medications or unfortunate recurrence due to medication failure.

We compared primary (SW480) and metastatic (SW620) cells via RNA sequencing and GO analysis, which demonstrated differential expression in cell adhesion and DNA repair mechanisms. The most remarkable difference is seen with angiogenesis factors between the two cell lines, with SW620 and SW480 demonstrating contrasting expression in VEGF and VEGFR1 (Figure 1E), once again pointing to another method in which cancer cells may transform and metastasize. It is also interesting to note the upregulation of EMT markers, such as vimentin and fibronectin, in epithelial cells that are usually considered to be more common with mesenchymal cells (Figure 1F). These findings correlate with our prior research that demonstrate fibronectin loss despite EMT of highly metastatic cells, pointing to another transformative framework [7] .

Western blots demonstrated little change in HR factor expression, while NHEJ repair mechanisms through Ku-80 are more prevalent in SW620. However, Ku-70 is maintained and approximately similarly expressed in SW620. SW620 demonstrated activation of DNA-PKcs, revealing a mechanism in which greater mutative effects may contribute to the metastatic transformation of CRC cells. SW620F appears to up-regulate NHEJ still further (Figure 2). This finding would also explain the sensitivity of SW620 and SW620F cells to the NHEJ inhibitor Nu-7441. Nu-7441 additionally showed differential response between primary and metastatic cell lines on an hourly basis seen with live microscopy. Metastatic cells likely rely more heavily on the NHEJ pathway, thus inhibiting NHEJ blocks their proliferation, a trend also seen in ovarian cancer cells [3] .

All three cell lines exhibited sensitivity to cediranib. Cediranib is a VEGF inhibitor and was seen to be comparable to bevacizumab, another VEGF inhibitor previously used in first line treatment of advanced metastatic CRC [10] . Investigation into the drug is ongoing, and findings correlate with the efficacy of using cediranib for metastatic CRC. Interestingly, Kaplan et al. showed that cediranib could also impact DNA repair, downregulating HR in cancer cells [11] .

Our RNA sequencing provides greater insight into DNA expression between primary CRC and metastatic CRC, revealing SW620 increases expression of angiogenesis, adhesion, and specifically genes such as PROM1 (prominin-1), a transmembrane protein expressed in adult stem cells that has been suggested as a prognostic marker in colorectal cancer [12] .

## 5 CONCLUSION

Our findings indicate that primary and metastatic colorectal cancer have different mechanisms for survival on a cellular level. Secondary (SW620) CRC demonstrates greater expression and activation of NHEJ factors compared to primary CRC, alluding to its potential to enhance its repair and survival. These findings may also point to a therapeutic opportunity - chemotherapeutic drugs need to be targeted towards cellular DNA repair mechanisms to target metastatic CRC, rather than using broad stroke chemotherapy to treat both primary and secondary CRC.

We found that cediranib and Nu-7441 show the greatest discrepancies between SW480 and SW620, and even greater when compared to SW620F. Drug response aligns with RNA sequencing and GO analysis. Whether the effects of cediranib are mediated by loss of DNA repair or angiogenic signals will be the subject of future studies. Overall, each of these drug responses provide insight into the ability of metastatic cells to improve their survival through DNA repair and angiogenesis.

## AUTHOR CONTRIBUTIONS

Josephine Tsang: Data curation, formal analysis, investigation, writing—original draft. Cai M. Roberts: Conceptualization, data curation, formal analysis, funding acquisition, methodology, resources, writing - review & editing.

## ETHICS STATEMENT

Not applicable.

## CONSENT

All authors critically revised the manuscript and consent for publication.

## DATA AVAILABILITY STATEMENT

RNA sequencing dataset is available on the NCBI GEO platform, accession number GSE308351. All our studies show that SW620F correlates with variable gene expression (increased NHEJ repair proteins) and drug resistance from its original line. The data generated and analyzed in the study are included in this manuscript.

## ACKNOWLEDGMENTS

The authors wish to thank Mr. Shanay Patel and Mr. Jihad Aburas for assistance with experiments. We thank Dr. Ellen Kohlmeir and the Midwestern University Core Facility, Downers Grove, IL. BCA assays were performed using the PerkinElmer EnSpire Plate Reader, and cells were counted using the Denovix CellDrop automated cell counter located in the Core. RNA sequencing was performed by Azenta Life Sciences (South Plainfield, NJ, USA).

